# Challenging the notion of a task-negative network: default mode network involvement under varying levels of cognitive effort

**DOI:** 10.1101/2021.11.03.466925

**Authors:** Sarah Weber, André Aleman, Kenneth Hugdahl

## Abstract

Everyday cognitive functioning is characterized by constant alternations between different modes of information processing, driven by fluctuations in environmental demands. At the neural level, this is realized through corresponding dynamic shifts in functional activation and network connectivity. A distinction is often made between the Default Mode Network (DMN) as a *task-negative* network that is upregulated in the absence of cognitive demands, and *task-positive* networks that are upregulated when cognitive demands such as attention and executive control are present. Such networks have been labelled the Extrinsic Mode Network (EMN). We investigated changes in brain activation and functional network connectivity during repeated alternations between levels of cognitive effort. Using fMRI and a block-design Stroop paradigm, participants switched back and forth between periods of no effort (resting), low effort (word reading, automatic processing) and high effort (color naming, cognitive control). Results showed expected EMN-activation for task versus rest, and likewise expected DMN-activation for rest versus task. The DMN was also more strongly activated during low effort contrasted with high effort, suggesting a gradual up- and down-regulation of the DMN, depending on the level of demand. The often reported “anti-correlation” between DMN and EMN was only present during periods of low effort, indicating intermittent contributions of both networks. These results challenge the traditional view of the DMN as solely a task-negative network. Instead, the present results suggest that both EMN and DMN may contribute to low-effort cognitive processing. In contrast, periods of resting and high effort are dominated by the DMN and EMN, respectively.

## 1. Introduction

The Default Mode Network (DMN) was discovered as a set of interconnected brain regions which are typically downregulated during the presence of external tasks or stimuli (Buckner et al., 2008; Raichle et al., 2001; Shulman et al., 1997). The DMN is therefore often labelled as a “task-negative” network, meaning that it is upregulated in the absence of demands for cognitive effort (Anticevic et al., 2012; Hugdahl et al., 2019). However, it is clear that sub-components, or nodes, of this network are present during cognitive processing, suggesting a more dynamic and complex role (Bressler & Menon, 2010; Greicius et al., 2003; Ossandón et al., 2011). This opens up the question how the DMN relates to what has been labelled “task-positive” networks (Uddin et al., 2009; Yamashita et al., 2020). Task-positive networks describe brain nodes and their functional connections that are upregulated in response to external stimulation and active task-processing. Apart from domain-specific networks, such as the visual or auditory networks, there are also non-specific task-positive networks which are activated across different cognitive domains (Duncan & Owen, 2000; Fedorenko et al., 2013; Hugdahl et al., 2015; Hugdahl et al., 2019; Riemer et al., 2020). Following the taxonomy introduced by Hugdahl et al. (2015), we will denote these task non-specific activations as an extrinsic mode network (EMN), in contrast to the DMN which is typically denoted as an intrinsic mode network (Fox et al., 2009; Raichle, 2010). The EMN has a morphological architecture with a fronto-temporo-parietal distribution, including the inferior and middle frontal gyri, inferior parietal lobule, supplementary motor area, and the inferior temporal gyrus (Hugdahl et al., 2015). It thus overlaps with what Fedorenko et al. (2013) called the cognitive flexibility network and Duncan called the multiple demand system. In addition, the EMN overlaps with domain-specific task-positive networks such as the salience, dorsal attention, and central executive networks (Bressler & Menon, 2010; Corbetta et al., 2008; Greicius et al., 2003). Although EMN nodes correlate negatively with DMN nodes (Hugdahl et al., 2019; Riemer et al., 2020), also called an anti-correlation (Fox et al., 2005; Provost & Monchi, 2015), there is evidence that the up- and down-regulation of task-negative and task-positive networks follows a gradual course when experimental demands abruptly alternate between rest and task-processing (Hugdahl et al., 2019; Riemer et al., 2020). These findings suggest that the relationship between the DMN as a task-negative and the EMN as a task-positive network is more flexible than previously thought, with dynamic changes in the relative contributions of the two networks to brain functioning (Elton & Gao, 2015; Gao et al., 2013; Piccoli et al., 2015). In everyday life, cognitive functioning is usually determined by frequent switching between different “tasks” with varying processing demands and with varying degrees of cognitive effort. Accordingly, brain functioning is likely to be characterized by a continuous shifting between varying contributions of task-negative and task-positive networks. For example, periods of highly effortful cognitive processing may engage the EMN more strongly than periods of lower cognitive effort (McKiernan et al., 2003). While DMN activity has traditionally been viewed to interfere with performance on cognitive tasks (Spreng, 2012), we suggest that the DMN may also contribute positively to performance under certain circumstances. Specifically, tasks that require low-effort, automatic processing seem to engage the DMN more than high-effort tasks. Provost et al. (2012, see also Provost & Monchi, 2015) used the Wisconsin Card Sorting task, with a repetitive, low-effort condition where subjects had to sort the cards according to the same rule over a longer period of time, and a high-effort condition where they had to switch between sorting rules. The results showed increased DMN activation in the former compared to the latter condition, suggesting a contribution of DMN to low-, but not high-effort cognitive processing. Similarly, Vatansever et al. (2017) divided performance on the Wisconsin Card Sorting task into a high-effort learning phase where participants had to work out the correct sorting rule, and a low-effort phase where they only had to apply this rule over several trials. DMN activity was increased in the first phase compared to the second phase of the task. Thus, it is reasonable to assume that the DMN contributes to externally driven, goal-directed cognition, and is up- and downregulated dynamically dependent on the type of processing degree of effort that is required at any point in time (cf. Riemer et al., 2020).

We aimed to follow-up these earlier attempts by investigating different degrees of cognitive effort, with a focus on interactions between the DMN and EMN networks by combining analyses of BOLD activation and functional connectivity. Cognitive effort was manipulated in an ON-OFF block-design based on the Stroop task (Stroop, 1935) with three experimental conditions. In a *high-effort* condition, participants named the ink color of the color words presented, irrespective of the words’ meaning. This condition is cognitively demanding since it requires active inhibition of the interfering meaning of the words, in addition to redirection of attention. In a *low-effort* condition, stimulus presentation was identical to the high-effort condition, but participants were instructed to read the color words irrespective of the color they were presented in. Since reading is an overlearned response, this condition is characterized by automatic, low-effort cognitive processing. In a third *no-effort* condition, participants passively viewed a fixation cross, with no words presented. We predicted that the EMN would be more activated during the high-effort ink-naming compared to the low-effort word reading condition, and during low-effort word reading compared to no-effort resting. Similarly, we predicted that the DMN would be more activated during the no-effort resting compared to the low-effort word reading, and during the low-effort word reading compared to the high-effort ink naming. With regard to functional connectivity, most previous research in this context has focused on comparisons between task and rest (e.g. Elton & Gao, 2015; Gao et al., 2013). We expected differential DMN connectivity when comparing high- and low effort conditions, with increased negative connectivity between then DMN and EMN under low-effort processing compared to high-effort processing (cf. Vatansever et al., 2017).

## 2. Methods

### 2.1 Participants

44 healthy adults were recruited through flyers in the university and university hospital environment. The mean age was 26.18 years (SD=4.37, range=21-38 years). 25 participants were female. All participants were right-handed native Norwegian speakers and reported normal color vision and no history of dyslexia or psychiatric or neurological conditions. Participants gave informed consent prior to participation. The study was conducted according to the Declaration of Helsinki and had received approval from the Norwegian ethics authorities (Regional Committees for Medical and Health Research Ethics (REK), project reference 2017/2452).

### 2.2 Materials and Procedure

#### 2.2.1 Stroop task

Participants performed a version of the Stroop paradigm (Stroop, 1935) that included three different levels of cognitive effort. In the high-effort condition, participants were asked to name the print color of the words that were shown, ignoring the meaning of the word. In the low-effort condition, participants were asked to read the color words out loud. An additional no-effort condition consisted of passive viewing of a fixation cross. Participants were instructed of the procedure for the three conditions and given examples before going into the MR scanner.

The three conditions were presented in alternating 30-second blocks (ABCACB…), each starting with a 3-second prompt screen that informed about which condition was to follow. There were eight blocks per condition. Each of the two task-blocks consisted of 18 stimuli presented in pseudorandom order with no more than three consecutive repetitions of identical stimuli, same color word or same printing color. The stimuli were four Norwegian color-words “blå” (blue), “grønn” (green), “gul” (yellow), and “rød” (red) written in blue, green, yellow or red ink in the center of the LCD screen, on black background. Presentation time was 250 ms with a 1500 ms gap between words in which participants gave their response verbally. Stimuli were presented with the E-prime software (version 2.0; https://pstnet.com/products/e-prime/) and viewed through MR-compatible LCD goggles which were adjusted to individual eyesight. Visibility was verified by each participant before the experiment started. Participants’ responses were recorded through an MR-compatible microphone and recording device which was attached to the head coil.

#### 2.2.2 MRI scanning parameters

Data were acquired at the Haukeland University Hospital in Bergen, Norway, on a Siemens Prisma 3T MR scanner. A structural T1-weighted scan was acquired with a seven-minute sequence, using the following parameters: repetition time (TR) = 1800 ms, echo time (TE) = 2.28 ms, flip angle = 8°, voxel size = 1 × 1 × 1 cm, field of view (FOV) = 256 × 256. During the Stroop task, a functional EPI-scan was acquired with the following parameters: TR = 2000 ms, TE = 30 ms, flip angle = 90°, voxel size = 3.6 × 3.6 × 3.6 cm, 34 slices (10% gap between slices), FOV = 1380 × 1380. The scan lasted approximately 15 minutes. In total, participants spent approximately 40 minutes in the scanner, including resting-state and MR spectroscopy sequences which is not reported in the current paper.

#### 2.2.3 fMRI preprocessing and analysis

EPI-data were preprocessed with the SPM-software (SPM12, https://www.fil.ion.ucl.ac.uk/spm/software/spm12/), using a standard preprocessing pipeline with realignment of functional images, coregistration with the T1 image, normalization into Montreal Neurological Institute (MNI) standard space, and smoothing with a Gaussian kernel of FWHM 6 mm. Parameters from the SPM motion correction procedure were analyzed in order to check that there was no excessive head motion. None of the participants had movements of greater than one voxel along any of the three axes.

Brain activity analyses were conducted in SPM with a classic block-design model containing rest-blocks as an implicit baseline, using block onsets convolved with a double-gamma hemodynamic response function to define the different conditions. Contrasts between the three conditions were calculated to reflect differences in cognitive effort: low effort (word reading) – no effort (rest condition); high effort (ink color naming) – no effort (rest condition), high effort (ink color naming) – low effort (word reading), plus the reverse of these contrasts. All results were FEW-corrected at p=.008 (Bonferroni-corrected for the six contrasts).

Functional connectivity analyses were conducted in the Conn-toolbox (version v.17.f http://www.nitrc.org/projects/conn). Preprocessed data first went through the Conn-toolbox default denoising pipeline, where motion parameters and their first derivatives as well as the BOLD-signal from white matter and cerebrospinal fluid masks (first five principle components for each) were regressed out. Linear detrending and a band-pass filter of 0.008-0.09 Hz was applied. Subsequently, seed-based functional connectivity analyses were conducted, and high-effort blocks were contrasted with low-effort blocks. Seed regions were derived from activation clusters from the block-activation analyses, resulting in a DMN seed and an EMN seed (see 3.2.2 below for further details). All results were corrected for multiple comparisons using Conn’s default FDR-cluster correction with a voxel-level p<.001 and cluster-level p=.025 (Bonferroni-corrected for the two seeds).

## 3. Results

### 3.1 Behavioral results: Stroop performance

A paired-sample t-test was performed to compare performance between the low- and high-effort conditions. Accuracy of responses was significantly lower in the high-effort condition compared to the low-effort condition (*M* = 96.42%, *SD* = 2.81, and *M* = 99.11, *SD* = 1.09, respectively, *t*(43) = 7.13, *p*<.001.

### 3.2 fMRI results

#### 3.2.1 fMRI activation for different levels of cognitive effort

Comparing the high- and low-effort tasks to the resting condition resulted in bilateral activation of frontal/precentral, parietal, occipital and superior temporal areas (Figure 1). Activations for these two contrasts showed however extensive overlap. Comparing high-effort to low-effort directly only resulted in small activation clusters, with stronger activity for the high-effort condition in the left superior parietal gyrus, left precentral/middle frontal gyrus and supplementary motor area (Figure 1). Details on cluster statistics for the three contrasts can be found in Supplementary Tables S1-S3.

**Figure 1.**
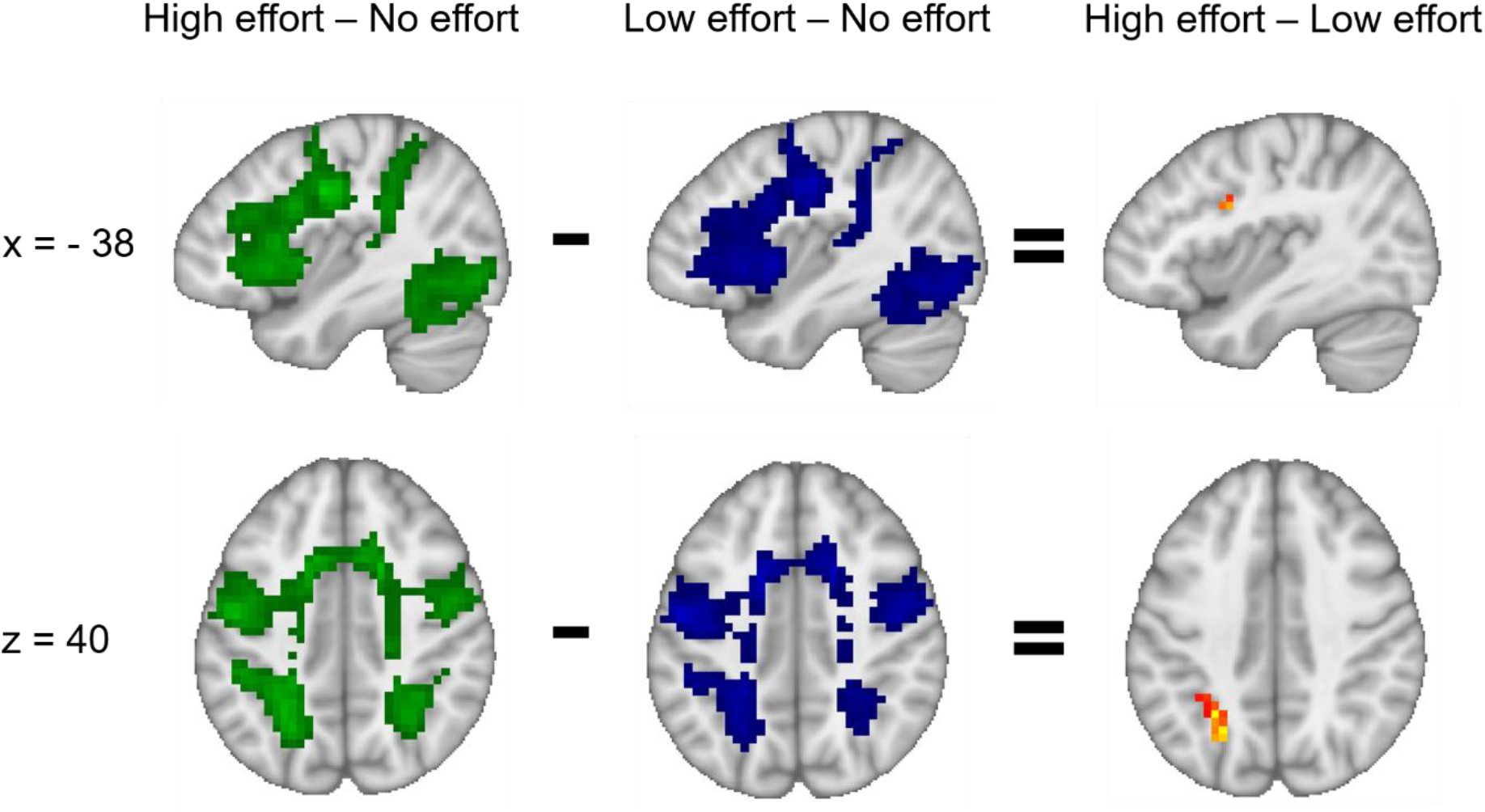
More intense activation in typical EMN areas for the comparisons of high effort versus rest (green), low effort versus rest (blue) and high effort versus low effort (yellow/red). The left side of the axial images corresponds to the left hemisphere.

Comparing the resting condition to high- and low-effort, respectively, resulted in significant activation in hub-areas of the DMN, in the inferior parietal cortices and in the anterior cingulate cortex (ACC) for both contrasts, and additionally in the posterior cingulate cortex (PCC) for the resting versus high-effort condition only (Figure 2). Comparing the two task-conditions with each other showed significantly higher activation in the parietal cortex, ACC and PCC for the low- versus high-effort contrast (Figure 2). Details on cluster statistics for the three contrasts can be found in Supplementary Tables S4-S6.

**Figure 2.**
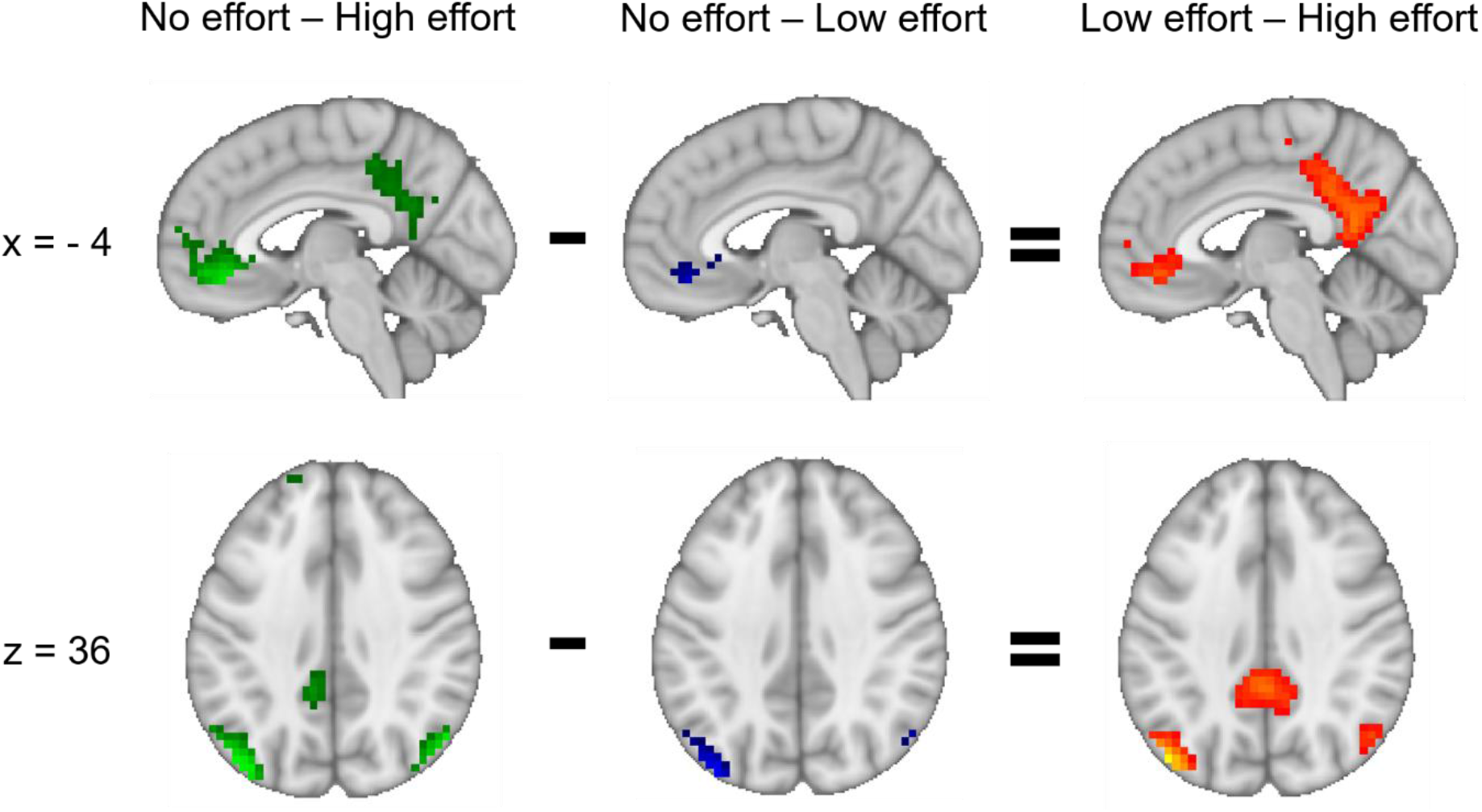
More intense activation in typical DMN areas for the comparisons of rest versus high effort (green), rest versus low effort (blue) and low effort versus high effort (yellow/red). The left side of the axial images corresponds to the left hemisphere.

#### 3.2.2 Functional connectivity for different levels of cognitive effort

The aim was to investigate functional connectivity between EMN and DMN areas that were susceptible to manipulations of cognitive effort. Therefore, activation clusters from the contrasts (high effort – low effort) as well as the (low effort – high effort) were used as seeds for the whole-brain connectivity analyses. This resulted in one seed comprising EMN areas (Figure 1, upper and lower right panel) and one seed comprising DMN areas (Figure 2, upper and lower right panel). For each of the two seeds, connectivity during low-effort blocks was compared to connectivity during high-effort blocks.

##### 3.2.2.1 EMN-seeded functional connectivity

During high effort compared to low effort, the EMN seed showed significantly stronger connectivity with other areas of the EMN, namely superior and middle frontal gyrus, pre-/postcentral gyrus, SMA and superior parietal cortex. Post-hoc analyses for low-effort and high-effort blocks separately showed positive connectivity during both conditions, but this positive connectivity was stronger during high-effort blocks. Furthermore, EMN-connectivity with the ACC and the PCC was significantly different during high effort compared to low effort, there was a significant negative connectivity between these regions during low-effort blocks which was not present during high-effort blocks (Figure 3). Statistics details for the EMN-seeded connectivity results can be found in Supplementary Table S7.

**Figure 3.**
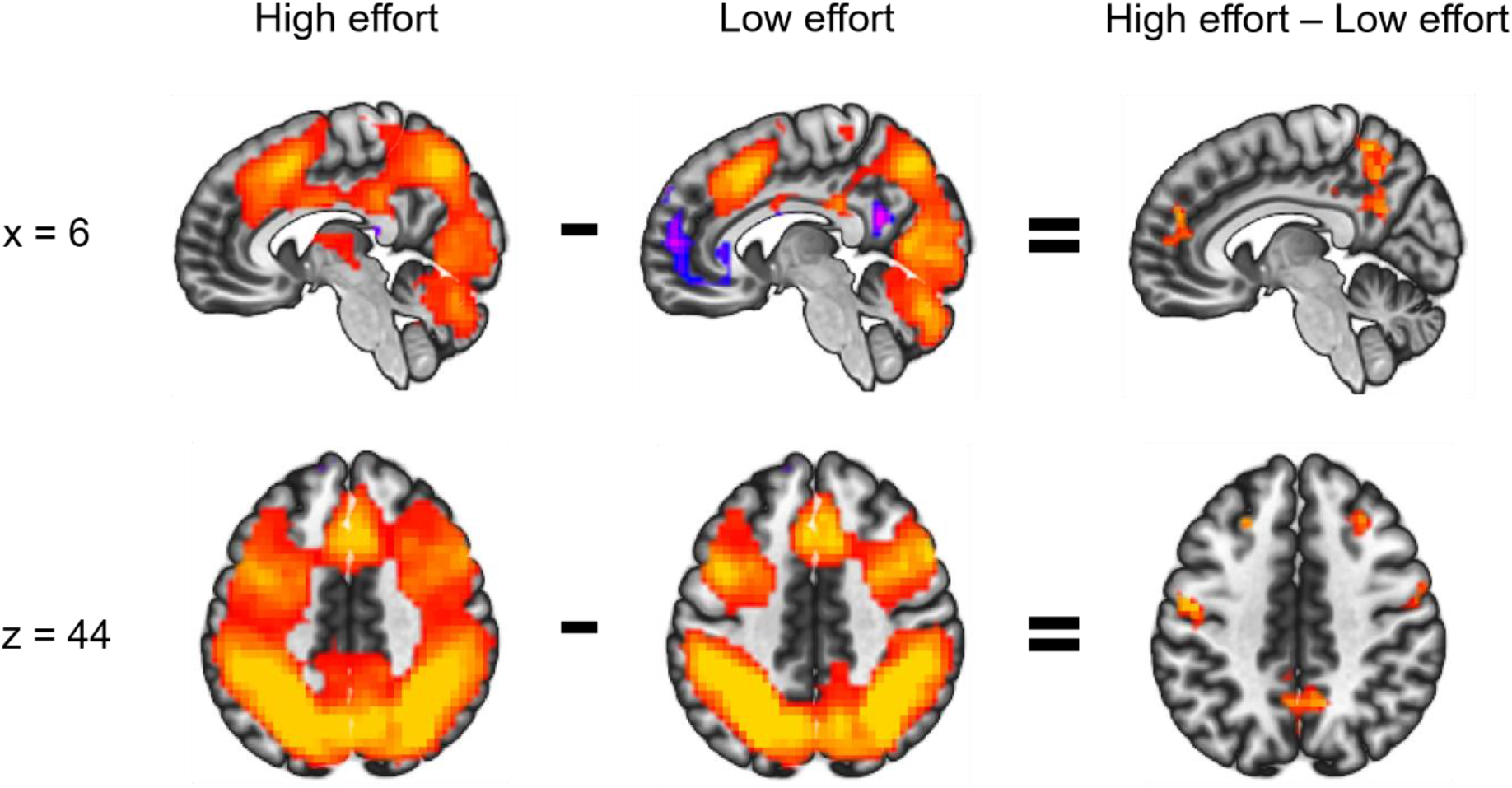
Sagittal and axial view of clusters that showed significant functional connectivity with the EMN seed (shown in Figure 1, right panel) during high effort (ink condition), during low effort (word condition), and differences when comparing the two conditions. Positive connectivity values are shown in red/orange and negative connectivity values in blue/purple. In the direct comparison, red indicates more positive connectivity values for high than low effort. The left side of the axial images corresponds to the left hemisphere.

##### 3.2.2.2 DMN-seeded functional connectivity

The DMN-seed showed significantly stronger connectivity with a small cluster in the PCC during low-effort compared to high-effort blocks. Connectivity was positive for both conditions separately but stronger for low-effort blocks. Furthermore, the DMN-seed showed significantly stronger negative connectivity with the SMA, superior and middle frontal gyrus, precentral gyrus, IFG and bilateral superior parietal cortex during low-effort compared to high-effort blocks. Post-hoc analyses showed that the difference in connectivity stemmed from a negative connectivity during low-effort blocks, not present during high-effort blocks (Figure 4). Statistics details on the DMN-seeded connectivity results can be found in Supplementary Table S8.

**Figure 4.**
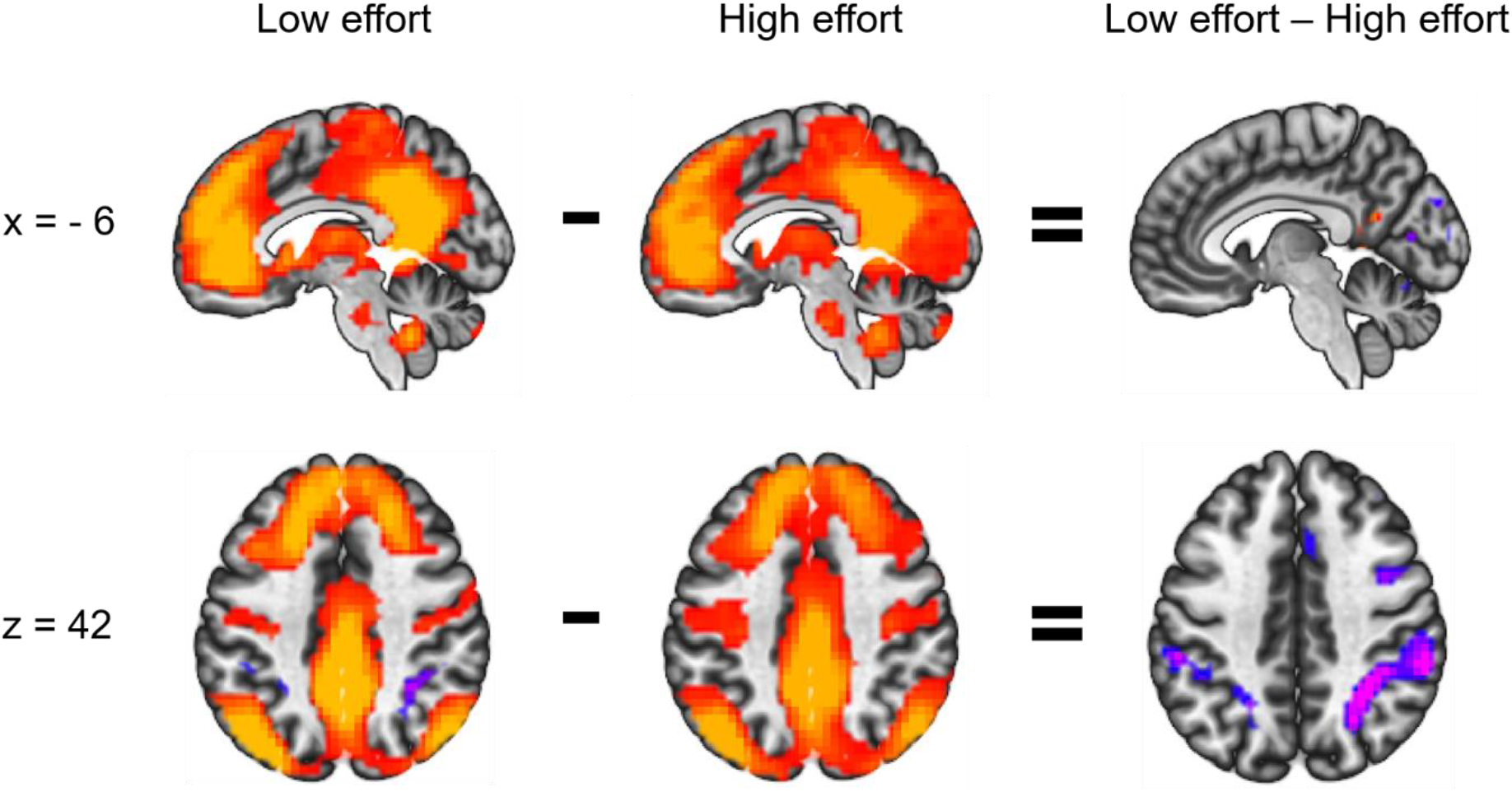
Sagittal and axial view of clusters that showed significant functional connectivity with the DMN seed (shown in Figure 2, right panel) during low effort (word condition), during high effort (ink condition), and differences when comparing the two conditions. Positive connectivity values are shown in red/orange and negative connectivity values in blue/purple. In the direct comparison, red indicates more positive connectivity values and blue more negative values for low effort than high effort. The left side of the axial images corresponds to the left hemisphere.

## 4. Discussion

The current study aimed to investigate two cortical networks, DMN and EMN, under conditions of varying cognitive effort. Functional activation as well as connectivity was examined while participants switched between processing task blocks with a high-effort condition that required cognitive control, a low-effort condition that required automatic processing, and a no-effort condition with passive resting. The degree of cognitive effort modulated activation in both networks, as well as connectivity between the networks. As expected, conditions of cognitive effort compared to rest activated fronto-parietal areas that have previously been found to be activated across a variety of cognitive tasks and together constitute the EMN (Hugdahl et al., 2015; see also Duncan, 2013; Fedorenko et al., 2013, Riemer et al., 2020). In line with previous research (Elton & Gao, 2015; Vatansever et al., 2017) frontal and parietal parts of the EMN were also significantly more activated during high effort compared to low effort processing, although the extent of these activation differences was relatively small. This finding suggests that the EMN is activated as soon as external stimulation or task demands are present, relatively independent of the level of effort involved, following a threshold mechanism of activation.

The areas that showed significantly more activation during resting compared with both task conditions have previously been described to constitute the “core DMN” (Buckner et al., 2008; Christoff et al., 2018). In line with recent findings (Vatansever et al., 2017), DMN areas in the present study were also more active during the low-effort compared to the high-effort condition, and this difference was even more pronounced than the difference between low-effort and resting. Thus, contrary to the traditional view of the DMN as a unique task-negative network which is down-regulated in the face of cognitive demands, or may even interferes with cognitive performance, these results indicate a more nuanced up- and downregulation depending on the amount of cognitive effort and load required. The largest difference between low and high effort was found in the PCC, which was downregulated only during high effort compared to resting, but not during low effort compared to resting. The PCC has often been described as a central hub for the DMN (Buckner et al., 2008; Christoff et al., 2018; Raichle, 2015). It may therefore be the “last man standing” when other nodes of the DMN are downregulated. Hence, the PCC could contribute to automatic processing, as in the low-effort condition. Taken together, these results suggest that up- and downregulation of the EMN and DMN is tuned in a way that the EMN is activated in a threshold-manner whenever there are requirements for controlled processing, even if the load is minimal, and additional activation or deactivation of the DMN is a regulatory parameter which is dependent on the degree of effort.

The areas of the DMN and the EMN that showed significant differential activation for low and high effort in the block-analysis, also showed differential effort-dependent connectivity patterns, when these areas were used as seed regions. The DMN showed stronger positive connectivity with its own core region, the PCC, during low effort compared to high effort. Increased connectivity between different DMN nodes has previously been found when comparing resting to cognitive processing (Elton & Gao, 2015; Gao et al., 2013), and when comparing low-effort to high-effort processing (Vatansever et al., 2017). Strong connectivity between DMN nodes has been interpreted as network stability and is positively correlated with cognitive performance (Mak et al., 2017) and negatively correlated with for example age (Grady et al., 2016). EMN-seeded connectivity was also modulated by cognitive effort. Comparing high- to low-effort processing, the EMN showed stronger positive connectivity with frontal and parietal areas which are typically associated with attention and cognitive control (Hugdahl et al., 2015; Passingham, 2021), thus showing increased within-connectivity between EMN nodes when task-demands and effort is high. Such increases could guide effortful cognitive processing, as suggested by previous findings of positive correlations between strong within-EMN connectivity and working memory load, as well as individual performance under conditions of high load (Liang et al., 2016).

Cognitive effort did not only modulate connectivity within networks but also connectivity between the two networks. The anti-correlation that has often been reported for task-negative and task-positive networks (Fox et al., 2005; Provost & Monchi, 2015) was significantly stronger during low-effort processing than during high-effort processing. In fact, post-hoc analyses for the two levels of cognitive effort separately showed that there was a significant anti-correlation between the DMN and EMN only for the low-effort condition. This was true when using both the DMN- and the EMN-seed, which supports the robustness of the finding. These results are in line with previously reported patterns of DMN connectivity (Vatansever et al., 2017), although these authors failed to find equivalent results when using a seed located in the task-positive network. A possible explanation for this discrepancy could be the exact location of the task-positive seed in the two studies. While Vatansever et al. (2017) used a frontal eye field ROI as a seed, a network-seed involving all areas with significantly greater activation for high compared to low effort was used in the present study. It seems that the whole network seed, rather than a single ROI seed, collectively contributes to differences in functional connectivity when levels of cognitive effort vary.

Overall, the present results indicate that both the EMN and DMN contribute to low-effort cognitive processing, in the sense that the EMN is upregulated during such periods, but the DMN is not fully downregulated. The anti-correlation between the two networks during these periods could suggest that their contributions happen intermittently. That is, periods of EMN dominance and DMN downregulation are alternated with states of DMN dominance and EMN downregulation, rather than a continuous parallel upregulation of both networks. In contrasts, periods of resting and periods of high effort are uniquely dominated by activation of the DMN and EMN, respectively. An interpretation of this is that under conditions of high cognitive effort activation in the two networks is independent of each other, which in turn could explain why there is no anti-correlation between the networks during such periods. We interpret this that under conditions of high to extreme cognitive effort, the task-positive EMN is so strongly up-regulated that it completely dominates and supersedes any dynamics related to the DMN. This is not the case during low-effort conditions, where both networks contribute, which in turn contributes to the anti-correlation between them. Whether the involvement of the DMN in low-effort conditions is task-related (and thus contributes to performance) or reflects independent activity remains to be further elucidated. Because our low-effort task concerned an automatized process (reading), performance was very high, precluding calculation of correlations with DMN involvement due to a ceiling effect. A study investigating different levels of working memory performance reported decreased engagement and responsiveness of the DMN in pain patients but performance was intact (i.e. not different from healthy subjects), without demonstrable compensatory neural recruitment (Čeko et al., 2016). The authors concluded that a responsive DMN might not be needed for successful cognitive performance.

The current results add to a growing body of evidence for an active role of the DMN in cognitive processing and a dynamic relationship with task-positive regions. However, as the present study has shown, the degree of such involvement is dependent on the level of cognitive effort that is required at any moment in time. In addition, the exact nature of the DMN’s contribution to cognition is still unclear. Sormaz et al. (2018) reported a contribution of the DMN to ongoing cognition, extending beyond task-unrelated processing, which is at odds with the task-negative view of the DMN. Using self-report data describing levels of detail of experience, relationship to a task, and emotional qualities they found that during periods of active working-memory maintenance, activity within the DMN was associated with the level of detail in ongoing thought. Other researchers have suggested that in particular the PCC as a highly connected core hub is involved in controlling cognition by monitoring changes in cognitive demands and integrating information from different networks in order to adjust behavior accordingly (Dohmatob et al., 2020; Leech et al., 2012). More recently, others have suggested that the PCC’s role is not only related to monitoring but also predicting cognitive demands in the environment (Araña-Oiarbide et al., 2020; Dohmatob et al., 2020). Such predictions would be easier during periods of highly rule-based automatic cognitive processing, which would explain greater engagement of the DMN during these periods.

In summary, we found that large-scale network dynamics during task-processing is dependent on the load of cognitive effort. During low-effort conditions, the DMN and EMN both contribute to the observed activation patterns, while activations in situations characterized by high effort and resting are separately driven by the EMN and DMN, respectively.

