## Supplementary tables for "Challenging the notion of a task-negative network: default mode network involvement under varying levels of cognitive effort"

### Supplementary material

Table S1
*Activation clusters for low cognitive effort versus no cognitive effort (word condition > rest condition)*

| | Size (k) | Sig. (p) | Peak (x y z) | t-value | Brain areas covered |
|---|---|---|---|---|---|
| Cluster 1 | 6207 | .000 | 13 -62 -20 | 18.37 | Cerebellum |
| | | .000 | 28 18 13 | 15.28 | AIns, FO |
| | | .000 | -44 -11 34 | 15.27 | PrG, PoG |

Peak locations are given in mm in MNI-152 standard space. Probabilistic locations are derived from the Neuromorphometrics, Inc. Atlas. Abbreviations: ACgG=anterior cingulate gyrus; AIns=anterior insula; AnG=angular gyrus; FO=frontal operculum; FRP=frontal pole; FuG=fusiform gyrus; Hipp=hippocampus; LiG=lingual gyrus; MFC=medial frontal cortex; MFG=middle frontal gyrus; MOG=middle occipital gyrus; MPrG=medial precentral gyrus; MSFG= medial superior frontal gyrus; PCu=precuneus; PCgG=posterior cingulate gyrus; PHG=parahippocampal gyrus; PoG=postcentral gyrus; PrG=precentral gyrus; SMC=supplementary motor cortex; SOG=superior occipital gyrus; SPL=superior parietal lobule

Table S2
*Activation clusters for high cognitive effort versus no cognitive effort (ink condition > rest condition)*

| | Size (k) | Sig. (p) | Peak (x y z) | t-value | Brain areas covered |
|---|---|---|---|---|---|
| Cluster 1 | 5986 | .000 | 13 -62 -16 | 18.98 | Cerebellum |
| | | .000 | -5 7 56 | 16.27 | SMC |
| | | .000 | -41 -11 34 | 15.92 | PrG, PoG |

Peak locations are given in mm in MNI-152 standard space. Probabilistic locations are derived from the Neuromorphometrics, Inc. Atlas. See Table S1 for abbreviations. Clusters smaller than k=10 are omitted for conciseness.

Table S3
*Activation clusters for high cognitive effort versus low cognitive effort (ink condition > word condition)*

| | Size (k) | Sig. (p) | Peak (x y z) | t-value | Brain areas covered |
|---|---|---|---|---|---|
| Cluster 1 | 32 | .000 | -26 -54 42 | 7.34 | SPL, AnG |
| Cluster 2 | 6 | .000 | -37 3 27 | 6.87 | PrG, MFG |

| | | | | | |
|---|---|---|---|---|---|
| Cluster 3 | 1 | .003 | -5 10 56 | 6.16 | SMC |

Peak locations are given in mm in MNI-152 standard space. Probabilistic locations are derived from the Neuromorphometrics, Inc. Atlas. See Table S1 for abbreviations. Clusters smaller than k=10 are omitted for conciseness.

Table S4
*Activation clusters for no cognitive effort versus low cognitive effort (rest condition > word condition)*

| | Size (k) | Sig. (p) | Peak (x y z) | t-value | Brain areas covered |
|---|---|---|---|---|---|
| Cluster 1 | 58 | .000 | -37 -80 38 | 9.64 | AnG, MOG |
| Cluster 2 | 61 | .000 | 6 32 -9 | 8.59 | ACgG, MFC, MSFG |
| Cluster 3 | 20 | .005 | 49 -72 27 | 8.06 | AnG, MOG |

Peak locations are given in mm in MNI-152 standard space. Probabilistic locations are derived from the Neuromorphometrics, Inc. Atlas. See Table S1 for abbreviations. Clusters smaller than k=10 are omitted for conciseness.

Table S5
*Activation clusters for no cognitive effort versus high cognitive effort (rest condition > ink condition)*

| | Size (k) | Sig. (p) | Peak (x y z) | t-value | Brain areas covered |
|---|---|---|---|---|---|
| Cluster 1 | 147 | .000 | -37 -80 38 | 12.64 | AnG |
| Cluster 2 | 65 | .000 | 53 -65 34 | 11.51 | AnG, MOG, SOG |
| Cluster 3 | 188 | .000 | 6 36 -9 | 11.34 | ACgG, MFC, MSFG, FRP |
| Cluster 4 | 146 | .000 | -16 -62 16 | 9.22 | PCu, PCgG, MPrG |
| Cluster 5 | 56 | .000 | 17 -58 24 | 8.51 | PCu, PCgG |
| Cluster 6 | 15 | .000 | -30 -36 -12 | 8.19 | FuG, Hipp, PHG, LiG |

Peak locations are given in mm in MNI-152 standard space. Probabilistic locations are derived from the Neuromorphometrics, Inc. Atlas. See Table S1 for abbreviations. Clusters smaller than k=10 are omitted for conciseness.

Table S6

*Activation clusters for low cognitive effort versus high cognitive effort (word condition > ink condition)*

| | Size (k) | Sig. (p) | Peak (x y z) | t-value | Brain areas covered |
|---|---|---|---|---|---|
| Cluster 1 | 88 | .000 | -44 -76 34 | 11.74 | AnG, MOG |
| Cluster 2 | 424 | .000 | 10 -51 13 | 9.06 | PCu, PCgG |
| Cluster 3 | 14 | .000 | -30 -36 -12 | 8.93 | FuG, Hipp, PHG |
| Cluster 4 | 88 | .000 | -1 43 -9 | 8.18 | ACgG, MFC, MSFG |
| Cluster 5 | 26 | .000 | 46 -69 38 | 7.86 | AnG, MOG |

Peak locations are given in mm in MNI-152 standard space. Probabilistic locations are derived from the Neuromorphometrics, Inc. Atlas. See Table S1 for abbreviations. Clusters smaller than k=10 are omitted for conciseness.

Table S7
*Clusters that show significantly different connectivity strength with the EMN seed during high cognitive effort versus low cognitive effort (ink condition > word condition)*

| | Size (k) | Sig. (p) | Peak (x y z) | Brain areas covered |
|---|---|---|---|---|
| Cluster 1 | 260 | .000 | 10 -58 56 | Precu, PCG |
| Cluster 2 | 95 | .000 | -52 -4 34 | PreCG, PostCG |
| Cluster 3 | 70 | .000 | 6 50 24 | aPaCiG, AC, FP, SFG |
| Cluster 4 | 49 | .000 | -19 14 -9 | Put, Caud, FOrb |
| Cluster 5 | 40 | .000 | 46 -11 56 | PreCG, PostCG |
| Cluster 6 | 28 | .003 | 13 -4 20 | Caud, Thal |
| Cluster 7 | 27 | .003 | 24 -36 52 | PostCG, SPL |

| | | | | |
|---|---|---|---|---|
| Cluster 8 | 27 | .003 | 24 28 42 | MidFG, SFG |
| Cluster 9 | 24 | .005 | 56 -65 20 | sLOC, AG, toMTG, iLOC |
| Cluster 10 | 22 | .006 | -16 -15 67 | SFG, SMA, PreCG |
| Cluster 11 | 21 | .007 | -26 28 42 | SFG, MidFG |

Peak locations are given in mm in MNI-152 standard space. Probabilistic locations are derived from the Harvard-Oxford Atlas. Abbreviations: AC=anterior cingulate gyrus; AG=angular gyrus; aPaCiG=anterior paracingulate gyrus; Caud=caudate; Cereb=cerebellum; CO=central opercular cortex Cun=cuneal cortex; FO=frontal operculum; FOrb=frontal orbital cortex; FP=frontal pole; IFG=inferior frontal gyrus; iLOC=inferior lateral occipital gyrus; Ins=insula; LG=lingual gyrus; MidFG=middle frontal gyrus; OFusG=occipital fusiform gyrus; OP=occipital pole; PCC=posterior cingulate gyrus; PO=parietal operculum; PostCG=postcentral gyrus; PreCG=precentral gyrus; Precu=precuneus; pSMG=posterior supramarginal gyrus; Put=putamen; SCC=supracalcarine cortex; SFG=superior frontal gyrus; SFG=superior frontal gyrus; sLOC=superior lateral occipital cortex; SMA=supplementary motor area; SPL=superior parietal lobule; Thal=thalamus; TOFusG=temporo-occipital fusiform gyrus; toITG=inferior temporo-occipital gyrus; toMTG=middle temporo-occipital gyrus

Table S8

*Clusters that show significantly different connectivity strength with the DMN seed during low cognitive effort versus high cognitive effort (word condition > ink condition)*

| | Size (k) | Sig. (p) | Peak (x y z) | Brain areas covered |
|---|---|---|---|---|
| Cluster 1 | 751 | .000 | 31 -47 42 | sLOC, pSMG, SPL, aSMG, ICC, OP, Cun, PostCG, SCC, AG, LG, PO, Precu |
| Cluster 2 | 524 | .000 | 53 18 -9 | FP, IFGoper, PreCG, MidFG, IC, FOrb, IFGtri |
| Cluster 3 | 127 | .000 | -55 -33 42 | aSMG, pSMG, PostCG, PO |
| Cluster 4 | 69 | .000 | -23 -58 45 | sLOC, SPL |
| Cluster 5 | 34 | .001 | -30 18 9 | |

| | | | | |
|---|---|---|---|---|
| | | | | Ins, FO, IFGoper, PreCG, CO |
| Cluster 6 | 29 | .003 | -30 -65 -5 | LG, OFusG, Cereb |
| Cluster 7 | 29 | .003 | -30 -69 -27 | Cereb, TOFusG, toITG |
| Cluster 8 | 28 | .003 | 2 10 56 | paCiG, SFG, SMA, AC |
| Cluster 9 | 27 | .003 | 60 -44 -2 | toMTG, toITG, pSMG |
| Cluster 10 | 26 | .003 | -26 -62 -45 | Cereb |
| Cluster 11 | 24 | .004 | 24 -65 -48 | Cereb |
| Cluster 12 | 19 | .012 | -5 -51 6 | PCC, Precu, |
| Cluster 13 | 16 | .022 | -44 -65 -2 | iLOC |
| Cluster 14 | 15 | .022 | -48 7 31 | PreCG, MidFG, IFGoper |
| Cluster 15 | 15 | .022 | -26 -69 24 | sLOC, Cun |
| Cluster 16 | 15 | .022 | -34 -44 -27 | Cereb, pTFusC, TOFusG |

Peak locations are given in mm in MNI-152 standard space. Probabilistic locations are derived from the Harvard-Oxford Atlas. See Table S7 for abbreviations.
